# Accelerating Wright-Fisher Forward Simulations on the Graphics Processing Unit

**DOI:** 10.1101/042622

**Authors:** David S. Lawrie

## Abstract

Forward Wright-Fisher simulations are powerful in their ability to model complex demography and selection scenarios, but suffer from slow execution on the CPU, thus limiting their usefulness. The single-locus Wright-Fisher forward algorithm is, however, exceedingly parallelizable, with many steps which are so-called *embarrassingly parallel*, consisting of a vast number of individual computations that are all independent of each other and thus capable of being performed concurrently. The rise of modern Graphics Processing Units (GPUs) and programming languages designed to leverage the inherent parallel nature of these processors have allowed researchers to dramatically speed up many programs that have such high arithmetic intensity and intrinsic concurrency. The presented <u>G</u>PU <u>O</u>ptimized Wright-<u>Fish</u>er simulation, or *GO Fish* for short, can be used to simulate arbitrary selection and demographic scenarios while running over 250-fold faster than its serial counterpart on the CPU. Even modest GPU hardware can achieve an impressive speedup of well over two orders of magnitude. With simulations so accelerated, one can not only do quick parametric bootstrapping of previously estimated parameters, but also use simulated results to calculate the likelihoods and summary statistics of demographic and selection models against real polymorphism data - all without restricting the demographic and selection scenarios that can be modeled or requiring approximations to the single-locus forward algorithm for efficiency. Further, as many of the parallel programming techniques used in this simulation can be applied to other computationally intensive algorithms important in population genetics, *GO Fish* serves as an exciting template for future research into accelerating computation in evolution. *GO Fish* is part of the Parallel PopGen Package available at: http://dl42.github.io/ParallelPopGen/

## Introduction

The Graphics Processing Unit (GPU) is commonplace in today’s consumer and workstation computers and provides the main computational throughput of the modern supercomputer. A GPU differs from a computer’s Central Processor Unit (CPU) in a number of key respects, but the most important differentiating factor is the number and type of computational units. While a CPU for a typical consumer laptop or desktop will contain anywhere from 2-4 very fast, complex cores, GPU cores are in contrast relatively slow and simple. However, there are typically hundreds to thousands of these slow and simple cores in a single GPU. Thus CPUs are low latency processors that excel at the serial execution of complex, branching algorithms. Conversely, the GPU architecture is designed to provide high computational bandwidth, capable of executing many arithmetic operations in parallel.

The historical driver for the development of GPUs was increasingly realistic computer graphics for computer games. However, researchers quickly latched on to their usefulness as tools for scientific computation – particularly for problems that were simply too time consuming on the CPU due to sheer number of operations that had to be computed, but where many of those operations could in principle be computed simultaneously. Eventually programming languages were developed to exploit GPUs as massive parallel processors and, overtime, the GPU hardware has likewise evolved to be more capable for both graphics and computational applications.

Population genetics analysis of single nucleotide polymorphisms (SNPs) is exceptionally amenable to acceleration on the GPU. Beyond the study of evolution itself, such analysis is a critical component of research in medical and conservation genetics, providing insight into the selective and mutational forces shaping the genome as well as the demographic history of a population. One of the most common analysis methods is the site frequency spectrum (SFS), a histogram where each bin is a count of how many mutations are at a given frequency in the population.

SFS analysis is based on the precepts of the Wright-Fisher process [1, 2], which describes the probabilistic trajectory of a mutation’s frequency in a population under a chosen evolutionary scenario. The defining characteristic of the Wright-Fisher process is forward time, non-overlapping, discrete generations with random genetic drift modeled as a binomial distribution dependent on the population size and the frequency of a mutation [1, 2]. On top of this foundation, can be added models for selection, migration between populations, mate choice & inbreeding, linkage between different loci, etc. For simple scenarios, an approximate analytical expression for the expected proportion of mutations at a given frequency in the population, the expected SFS, can be derived [1-5]. This expectation can then be compared to the observed SFS of real data, allowing for parameter fitting and model testing [5-7]. However, more complex scenarios do not have tractable analytical solutions, approximate or otherwise. One approach is to simulate the Wright-Fisher process forwards in time to build the expected frequency distribution or other population genetic summary statistics [8-11]. Because of the flexibility inherent in its construction, the Wright-Fisher forward simulation can be used to model any arbitrarily complex demographic and selection scenario [8-13]. Unfortunately, because of the computational cost, the use of such simulations to analyze polymorphism data is often prohibitively expensive in practice [12, 13]. The coalescent looking backwards in time and approximations to the forward single-locus Wright-Fisher algorithm using diffusion equations provide alternative, computationally efficient methods of modeling polymorphism data [14, 15]. However, these effectively limit the selection and demographic models that can be ascertained and approximate the Wright-Fisher forward process [12, 13, 15, 16]. Thus by speeding up forward simulations, we can use more complex and realistic demographic and selection models to analyze within-species polymorphism data.

Single-locus Wright-Fisher simulations based on the Poisson Random Field model [4] ignore linkage between sites and simulate large numbers of individual mutation frequency trajectories forwards in time to construct the expected SFS. Exploiting the naturally parallelizable nature of the single-locus Wright-Fisher algorithm, these forward simulations can be greatly accelerated on the GPU. Written in the programming language CUDA v6.5 [17], a C/C++ derivative for NVIDIA GPUs, the <u>G</u>PU <u>O</u>ptimized Wright-<u>Fish</u>er simulation, *GO Fish*, allows for accurate, flexible simulations of SFS at speeds orders of magnitude faster than comparative serial programs on the CPU. *GO Fish* can be both run as a standalone executable and integrated into other programs as a library to accelerate single-locus Wright-Fisher simulations used by those tools.

## Algorithm

In a single-locus Wright-Fisher simulation, a population of individuals can be represented by the set of mutations segregating in that population – specifically by the frequencies of the mutant, derived alleles in the population. Under the Poisson Random Field model, these mutations are completely independent of each other and new mutational events only occur at non-segregating sites in the genome (i.e. no multiple hits) [4].

Figure 1 sketches the algorithm for a typical, serial Wright-Fisher simulation, starting with the initialization of an array of mutation frequencies. From one discrete generation time step to the next, the change in any given mutation’s frequency is dependent on the strength of selection on that mutation, migration from other populations, the percent of inbreeding, and genetic drift. Unlike the others listed, inbreeding is not directly a force for allele frequency change, but rather it modifies the effectiveness of selection and drift. Frequencies of 0 (lost) and 1 (fixed) are absorbing boundaries such that if a mutation becomes fixed or lost across all extant populations, it is removed from the next generation’s mutation array. New mutations arising stochastically throughout the genome are then added to the mutation array of the offspring generation, replacing those mutations lost and fixed by selection and drift. As the offspring become the parents of the next generation, the cycle repeats until the final generation of the simulation.

**Figure 1.**
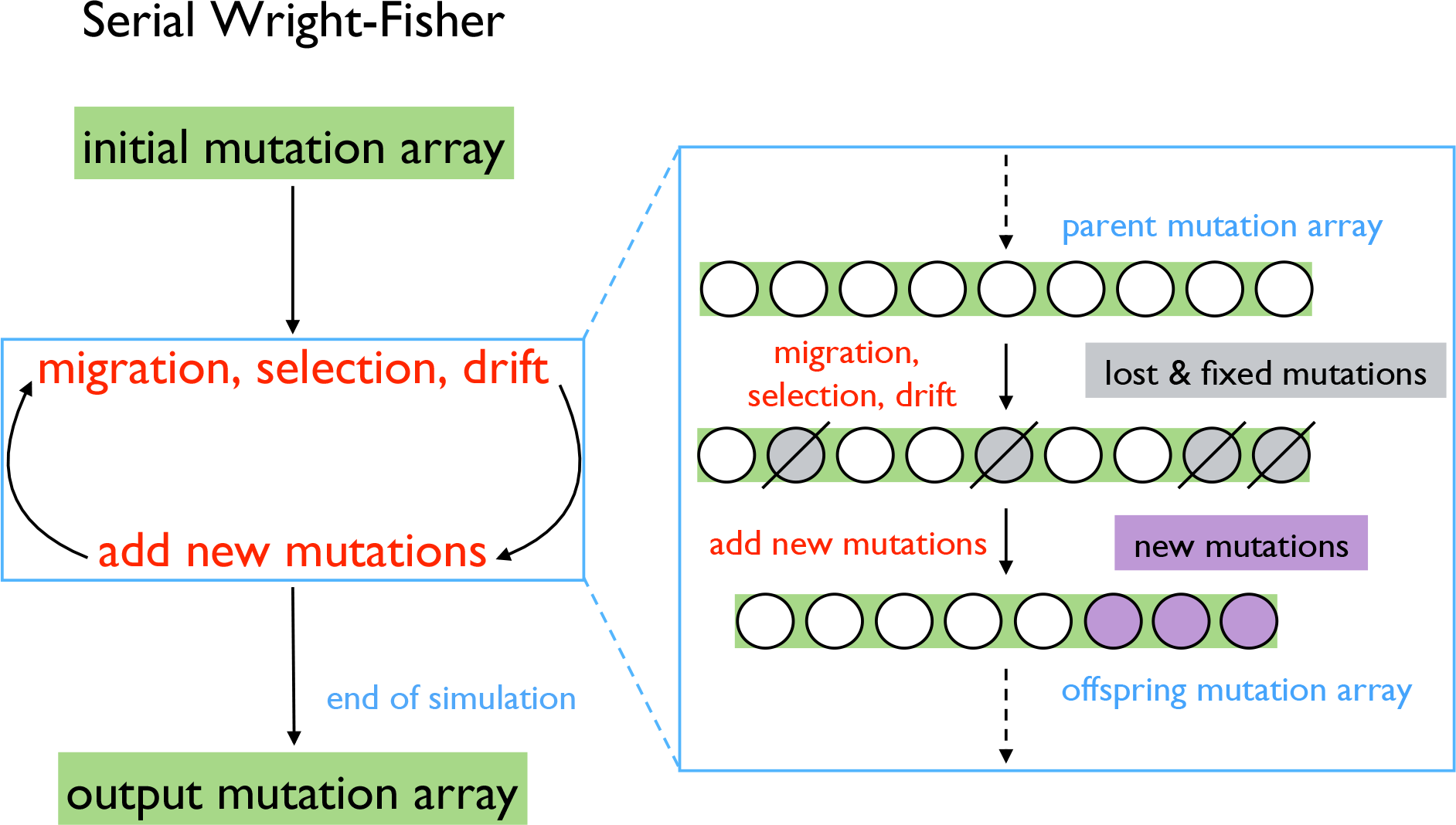
Serial Wright-Fisher algorithm. Mutations are the “unit” of simulation for the singlelocus Wright-Fisher algorithm. Thus a generation of organisms is represented by an array of mutations and their frequency in the (each) population (if there are multiple in the simulation). There are several options for how to initialize the mutation array to start a simulation: a blank mutation array, the output of a previous simulation run, or mutation-selection equilibrium (for details, see Appendix: *Simulation Initialization*). Simulating each discrete generation consists first of calculating the new allele frequency of each mutation, one at a time, where those mutations that become lost or fixed are discarded. Next, new mutations are added to the array, again, one at a time. The resulting offspring array of mutation frequencies becomes the parent array of the next generation and the cycle is repeated until the end of the simulation when the final mutation array is output.

While the details of how a GPU organizes computational work are quite intricate [17], the vastly oversimplified version is that a serial set of operations is called a thread and the GPU can execute many such threads in parallel. With completely unlinked sites, every simulated mutation frequency trajectory is independent of every other mutation frequency trajectory in the simulation. Therefore, the single-locus Wright-Fisher algorithm is trivially parallelized by simply assigning a thread to each mutation in the mutation array: when simulating each discrete generation, both calculating the new frequency of alleles in the next generation and adding new mutations to next generation are *embarrassingly parallel* operations (Figure 2A). This is the parallel ideal because no communication across threads is required to make these calculations. A serial algorithm has to calculate the new frequency of each mutation one by one – and the problem is multiplied where there are multiple populations, as these new frequencies have to be calculated for each population. For example, in a simulation with 100,000 mutations in a given generation and 3 populations, 300,000 sequential passes through the functions governing migration, selection, and drift are required. However, in the parallel version, this huge number of iterations can theoretically be compressed to a single step in which all the new frequencies for all mutations are computed simultaneously. Similarly, if there are 5,000 new mutations in a generation, a serial algorithm has to add each of those 5,000 new mutations one at a time to the simulation. The parallel algorithm can, in theory, add them all at once. Of course, a GPU only has a finite number of computational resources to apply to a problem and thus this ideal of executing all processes in a single time step is never truly realizable for a problem of any substantial size. Even so, parallelizing migration, selection, drift, and mutation on the GPU results in dramatic speedups relative to performing those same operations serially on the CPU. This is the main source of *GO Fish*’s improvement over serial, CPU-based Wright-Fisher simulations.

**Figure 2.**
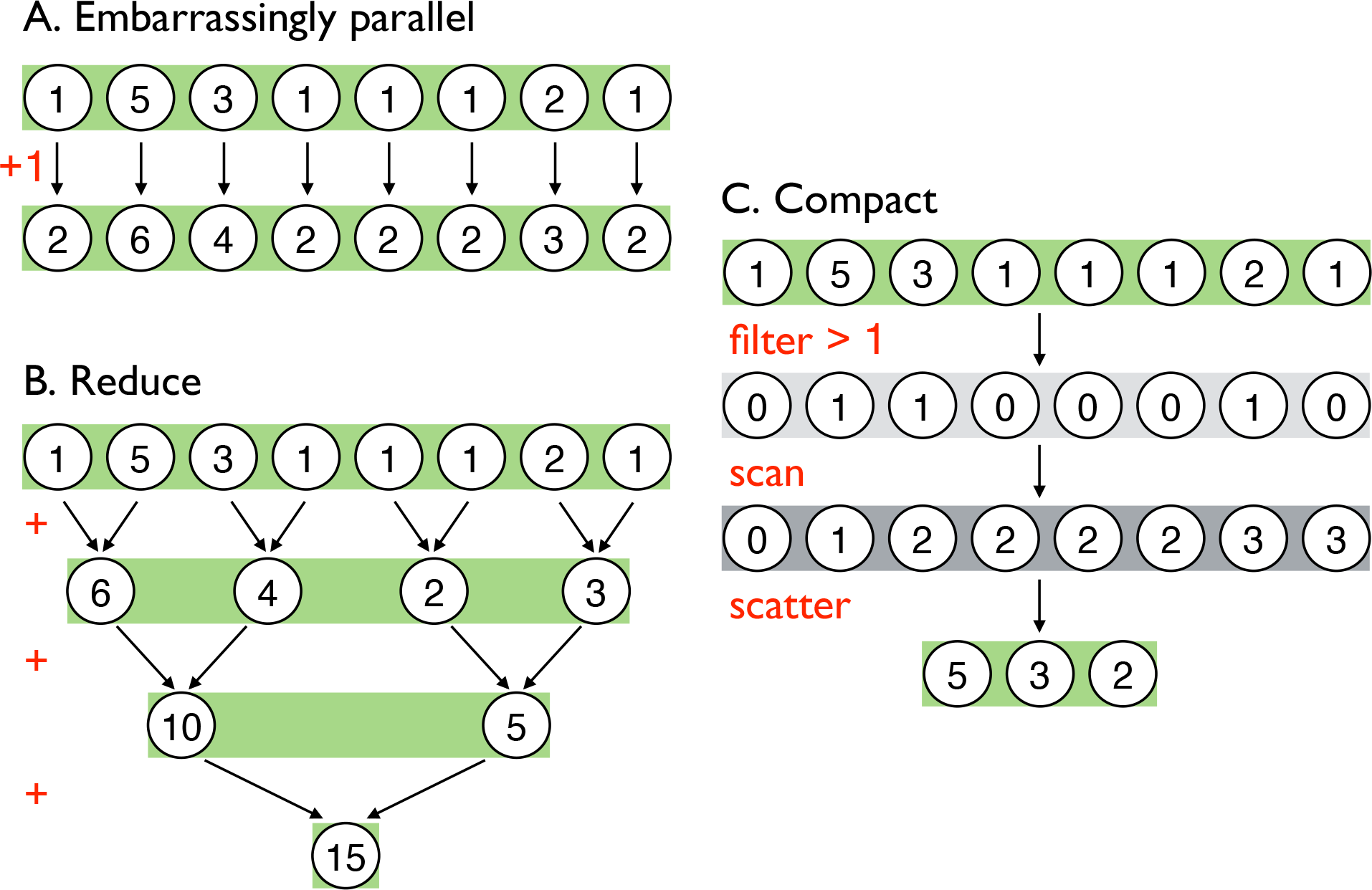
Common Parallel Algorithms. Above are illustrative examples of three classes of common parallel algorithms implemented using simple operations and an eight-element, integer array. **A) Embarrassingly parallel** algorithms are those that can be computed independently and thus simultaneously on the GPU. The given example, adding 1 to every element of an array, can be done concurrently to all array elements. In *GO Fish*, calculating new mutation frequencies and adding new mutations to population are both *embarrassingly parallel* operations. **B) Reduce** is a fundamental parallel algorithm in which all the elements of an array are reduced to a single value using a binary operator, such as in the above summation over the example array [55]. This algorithm takes advantage of the fact that in each time step half of the sums can be done independently while synchronized communication is necessary to combine the results of previous time steps. In total, log2(8) = 3 time steps are required to reduce the example array. **C) Compact** is a multi-step algorithm that allows one to filter arrays on the GPU [18]. In an *embarrassingly parallel* step, the algorithm presented above first creates a new Boolean array of those elements that passed the filter predicate (e.g. x > 1). Then a *scan* is performed on the Boolean array. *Scan* is similar in concept to *reduce*, wherein for each time step half of the binary operations are independent, but it is a more complex parallel algorithm that creates a running sum over the array rather than condensing the array to a single value (see [56]). This running sum determines the new index of each element in the original array being filtered and the size of the new array. Those elements that passed the filter are then scattered to their new indices of the now smaller, filtered array. *Compact* is used in *GO Fish* to filter out fixed and lost mutations.

One challenge to the parallelization of the Wright-Fisher algorithm is the treatment of mutations that become fixed or lost. When a mutation reaches a frequency of 0 (in all populations, if multiple) or 1 (in all populations, if multiple), that mutation is forever lost or fixed. Such mutations are no longer of interest to maintain in memory or process from one generation to the next. Without removing lost and fixed mutations from the simulation, the number of mutations being stored and processed would simply continue to grow as new mutations are added each generation. When processing mutations one at a time in the serial algorithm, removing mutations that become lost or fixed is as trivial as simply not adding them to the next generation and shortening the mutation array in the next generation by 1 each time. This becomes more difficult when processing mutations in parallel. As stated before: the different threads for different mutations do not communicate with each other when calculating the new mutation frequencies simultaneously. Therefore any given mutation/thread has no knowledge of how many other mutations have become lost or fixed that generation. This in turn means that when attempting to remove lost and fixed mutations while processing mutations in parallel, there is no way to determine the size of the next generation’s mutation array or where in the offspring array each mutation should be placed.

One solution to the above problems is the algorithm *compact* [18], which can filter out lost and fixed mutations while still taking advantage of the parallel nature of GPUs (Figure 2C). However, compaction is not *embarrassingly parallel*, as communication between the different threads for different mutations is required, and it involves a lot of moving elements around in GPU memory rather than intensive computation. Thus, it is a less efficient use of the GPU as compared to calculating allele frequencies. As such, a nuance in optimizing *GO Fish* is how frequently to remove lost and fixed mutations from the active simulation. Despite the fact that computation on such mutations is wasted, calculating new allele frequencies is so fast that not filtering out lost and fixed mutations every generation and temporarily leaving them in the simulation actually results in faster runtimes. Eventually of course, the sheer number of lost and fixed mutations overwhelms even the GPU’s computational bandwidth and they must be removed. How often to compact for optimal simulation speed can be ascertained heuristically and is dependent on the number of mutations each generation in the simulation and the attributes of the GPU the simulation is running on. Figure 3 illustrates the algorithm for *GO Fish*, which combines parallel implementations of migration, selection, drift, and mutation with a compacting step run every X generations and again before the end of the simulation.

**Figure 3.**
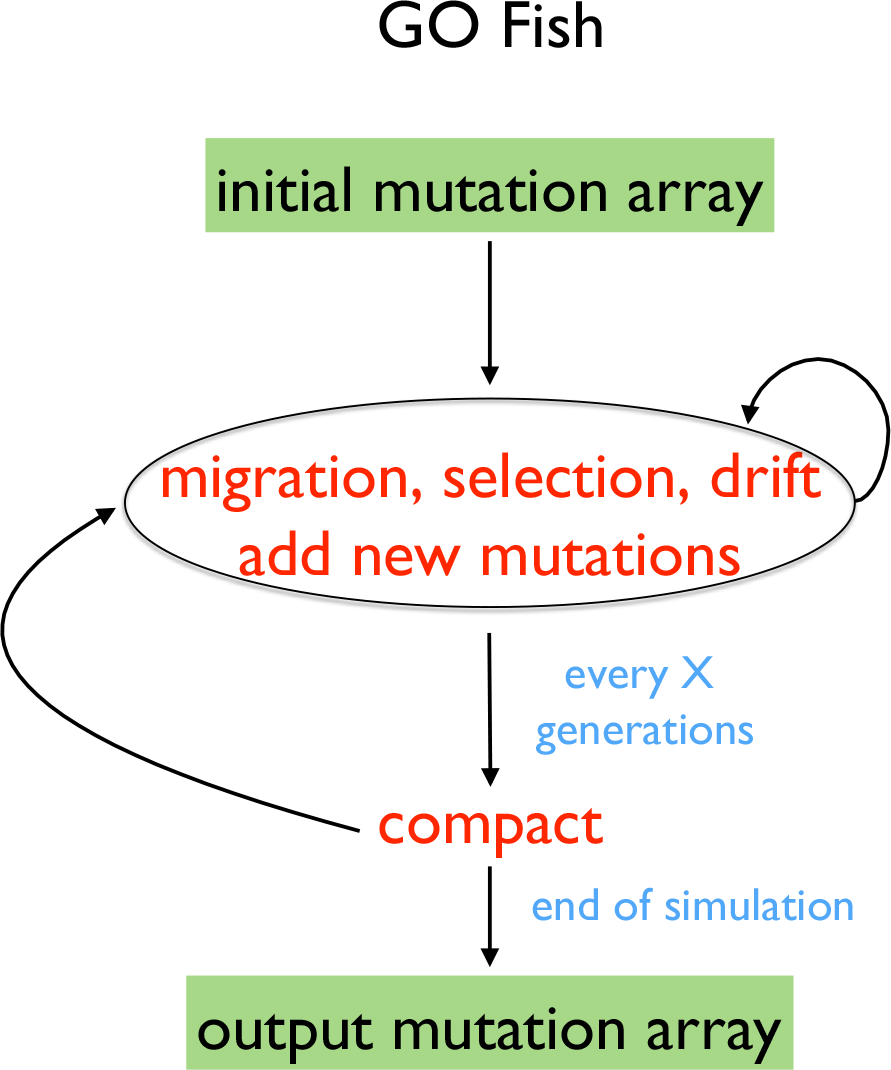
*GO Fish* algorithm. Both altering the allele frequencies of mutations from parent to child generation and adding new mutations to the child generation are *embarrassingly parallel* operations (see Figure 2A) that are greatly accelerated on the GPU. Further, as independent operations, adding new mutations and altering allele frequencies can be done concurrently on the GPU. In comparison to serial Wright-Fisher simulations (Figure 1), *GO Fish* includes an extra *compact* step (see Figure 2C) to remove fixed and lost mutations every X generations. Until compaction, the size of the mutation array grows by the number of new mutations added each generation. Before the simulation ends, the program compacts the mutation array one final time.

### The Population Genetics Model of GO Fish

A more detailed description of the implementation of the Wright-Fisher algorithm underlying *GO Fish*, with derivations of the equations below, can be found in the Appendix. Table 1 provides a glossary of the variables used in the simulation.

**Table 1.**
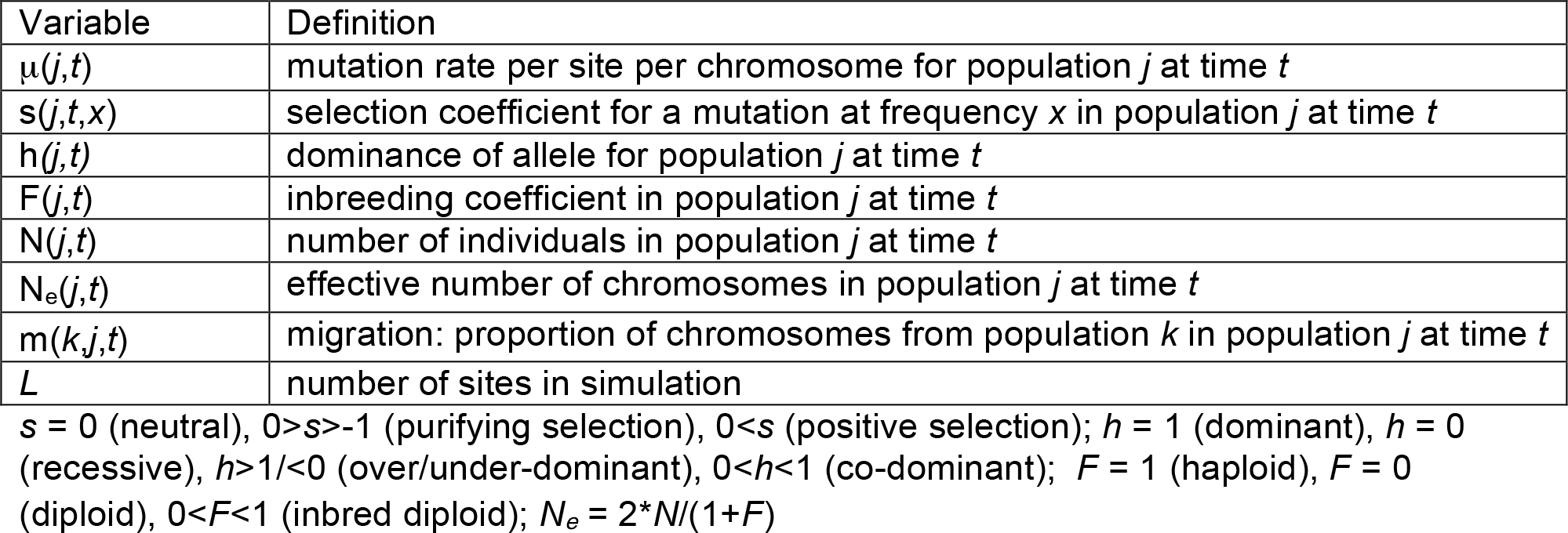
Glossary of simulation terms

| Variable | Definition |
| --- | --- |
| $\mu(j,t)$ | mutation rate per site per chromosome for population $j$ at time $t$ |
| $s(j,t,x)$ | selection coefficient for a mutation at frequency $x$ in population $j$ at time $t$ |
| $h(j,t)$ | dominance of allele for population $j$ at time $t$ |
| $F(j,t)$ | inbreeding coefficient in population $j$ at time $t$ |
| $N(j,t)$ | number of individuals in population $j$ at time $t$ |
| $N_e(j,t)$ | effective number of chromosomes in population $j$ at time $t$ |
| $m(k,j,t)$ | migration: proportion of chromosomes from population $k$ in population $j$ at time $t$ |
| $L$ | number of sites in simulation |

$s = 0$ (neutral), $0 > s > -1$ (purifying selection), $0 < s$ (positive selection); $h = 1$ (dominant), $h = 0$ (recessive), $h > 1/2 < 1$ (over/under-dominant), $0 < h < 1/2$ (co-dominant); $F = 1$ (haploid), $F = 0$ (diploid), $0 < F < 1$ (inbred diploid); $N_e = 2 \cdot N / (1 + F)$

The simulation can start with an empty initial mutation array, with the output of a previous simulation run, or with the frequencies of the initial mutation array in mutation-selection equilibrium. Starting a simulation as a blank canvas provides the most flexibility in the starting evolutionary scenario. However, to reach an equilibrium start point requires a “burn-in”, which may be quite a large number of generations [11]. To save time, if a starting scenario is shared across multiple simulations, then the post-burn-in mutation array can be simulated beforehand, stored, and input as the initial mutation array for the next set of simulations. Alternatively, the simulation can be initialized in a calculable, approximate mutation-selection equilibrium state, allowing the simulation of the evolutionary scenario of interest to begin essentially immediately. λ_μ_(*x*) is the expected (mean) number of mutations at a given frequency, *x*, in the population at mutation-selection equilibrium and can be calculate via the following equation:

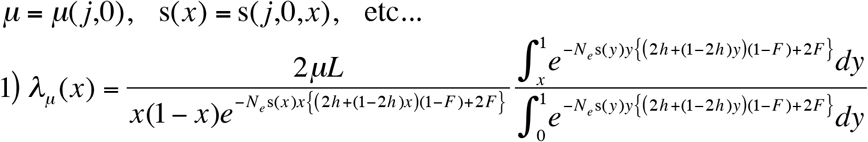

The derivation for eq. 1 can be found in the Appendix (eq. 1-6 in the Appendix). The numerical integration required to calculate λ_μ_(*x*) has been parallelized and accelerated on the GPU. To start the simulation, the actual number of mutations at each frequency is determined by draws from the Inverse Poisson distribution with mean and variance λ_μ_(*x*). This numerical initialization routine can handle most of the equilibrium evolutionary scenarios the main simulation is capable of itself – a major exception being those cases with migration between multiple populations. Given the number of cases covered by the above integration technique, this is likely to be the primary method to start a *GO Fish* simulation in a state of mutationselection equilibrium.

After initialization begins the cycle of adding new mutations to the population and calculating new frequencies for currently segregating mutations. The number of new mutations introduced in each population *j*, for each generation *t* is Poisson distributed with mean *N_e_μL* in accordance with the assumptions of the Poisson Random Field Model. These new mutations start at frequency 1/*N_e_* in the simulation. Meanwhile, the SNP frequencies of the extant mutations in the current generation *t*, and population *j* are modified by the forces of migration (I.), selection (II.), and drift (III.) to produce the new frequencies of those mutations in generation *t+1*.

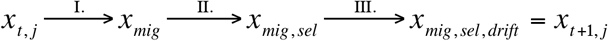

I. *GO Fish* uses a conservative model of migration [19] where the new allele frequency, *xmig*, in population *j* is the average of the allele frequency in all the populations weighted by the migration rate from each population, to population *j*. II. Selection further modifies the expected frequency of the mutations in population *j* according to eq. 2 below:

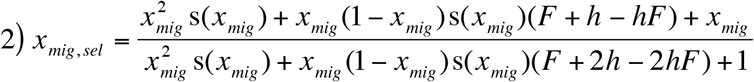

The derivation for eq. 2 can be found in the Appendix (eq. 8-13 in the Appendix). The variable *X_mig,sel_* represents the expected frequency of an allele in generation *t+1*. III. Drift, which is modeled as a binomial random deviation with mean *N_e_x_mig,sel_* and variance *N_e_x_mig,sel_*(1-*X_mig,sel_*), then acts on top of the deterministic forces of migration and selection to produce the ultimate frequency of the allele in the next generation, *t+1*, in population *j, X_t+1,j_*. Then the cycle repeats.

## Results and Discussion

To test the speed improvements from parallelizing the Wright-Fisher algorithm, *GO Fish* was compared to a serial Wright-Fisher simulation written in C++. Each program was run on two computers: an iMac and a self-built Linux-box with equivalent Intel Haswell CPUs, but very different NVIDIA GPUs. Constrained by the thermal and space requirements of laptops and all-in-one machines, the iMac’s NVIDIA 780M GPU (1536 cores@823 MHz) is slower and older than the NVIDIA 980 (2048 cores@1380MHz) in the Linux-box. For a given number of simulated populations and number of generations, a key driver of execution time is the number of mutations in the simulation. Thus many different evolutionary scenarios will have similar runtimes if they result in similar numbers of mutations being simulated each generation. As such, to benchmark the acceleration provided by parallelization and GPUs, the programs were run using a basic evolutionary scenario while varying the number of expected mutations in the simulation. The utilized scenario is a simple, neutral simulation, starting in mutation-selection equilibrium, of a single, haploid population with a constant population size of 200,000 individuals over 1,000 generations and a mutation rate of 1x10-9 mutations per generation per individual per site. With these other parameters held constant, varying the number of sites in the simulation adjusts the number of expected mutations for each of the benchmark simulations.

As shown in Figure 4: accelerating the Wright-Fisher simulation on a GPU results in massive performance gains on both an older, mobile GPU like the NVIDIA 780M and a newer, desktop-class NVIDIA 980 GPU. For example, when simulating the frequency trajectories of ∽500,000 mutations over 1,000 generations, *GO Fish* takes ∽0.2s to run on a 780M as compared to ∽18s for its serial counterpart running on the Intel i5/i7 CPU (@3.9 Ghz), a speedup of 88-fold. On a full, modern desktop GPU like the 980, *GO Fish* runs this scenario ∽176x faster than the strictly serial simulation and only takes about 0.1s to run. As the number of mutations in the simulation grows, more work is tasked to the GPU and the relative speedup of GPU to CPU increases logarithmically. Eventually though, the sheer number of simulated SNPs saturates even the computational throughput of the GPUs, producing linear increases in runtime for increasing SNP counts, like for serial code. Thus, eventually, there is a flattening of the fold performance gains. This plateau occurs earlier for 780M than for the more powerful 980 with its more and faster cores. Executed serially on the CPU, a huge simulation of ∽4x10^7^ SNPs takes roughly 24min to run versus only ∽13s/5.7s for *GO Fish* on the 780M/980, an acceleration of more than 109/250-fold. While not benchmarked here, the parallel Wright-Fisher algorithm is also trivial to partition over multi-GPU setups in order to further accelerate simulations.

**Figure 4.**
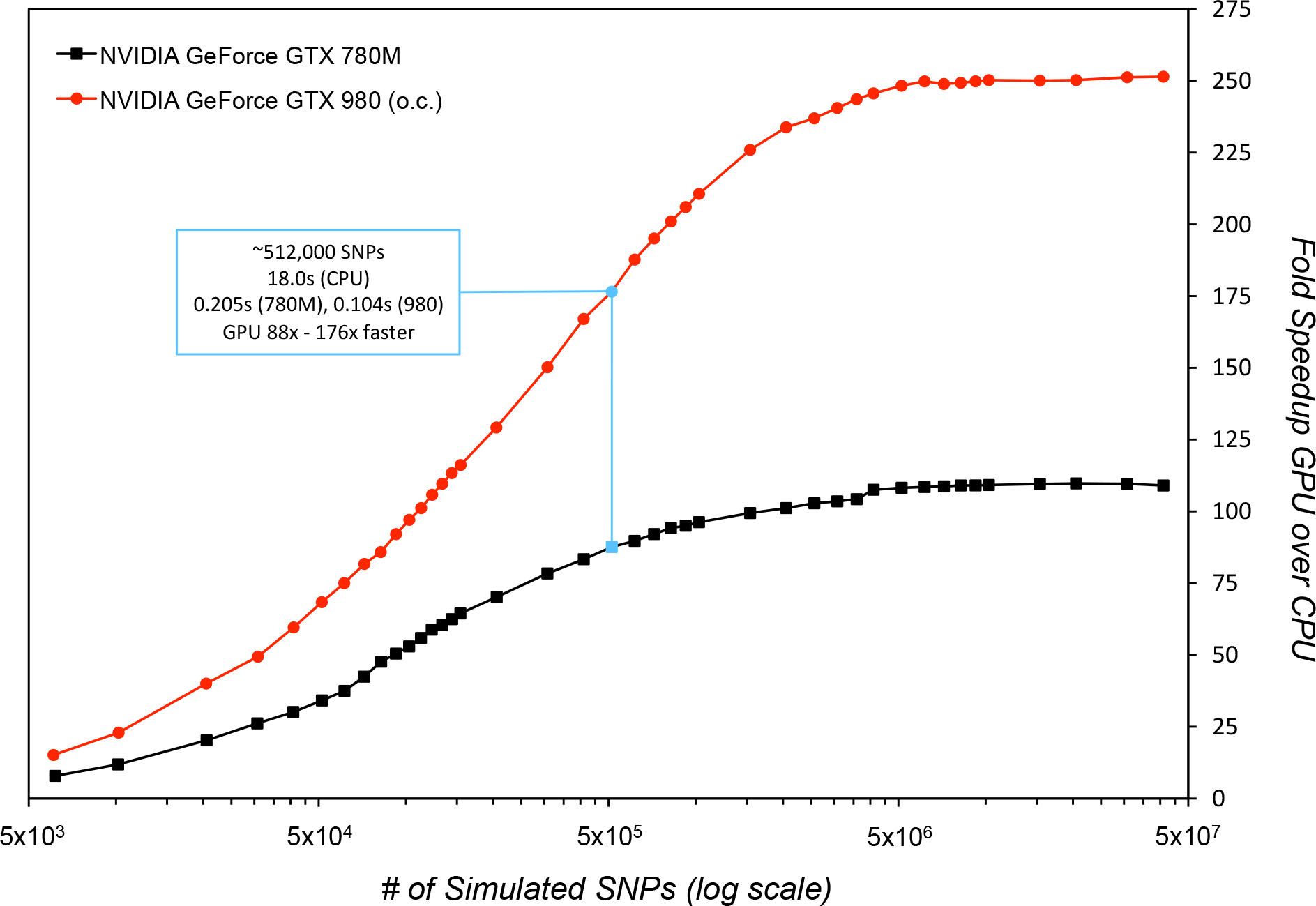
Performance gains on GPU relative to CPU. The above figure plots the relative performance of *GO Fish*, written in CUDA, to a basic, serial Wright-Fisher simulation written in C++. The two programs were run both on a 2013 iMac with an NVIDIA GeForce GTX 780M mobile GPU, 1536@823 MHz cores, (black line) and an Intel Core i7 4771 <u>CPU@3.9</u> GHz and a self-built Linux-box with a factory-overclocked NVIDIA GeForce GTX 980 GPU, 2048@1380MHz cores, and an Intel Core i5 4690K <u>CPU@3.9GHz</u> (red line). Full compiler optimizations (-O3 –fast-math) were applied to both serial and parallel programs. Each dot represents a simulation run plotted by the number of SNPs in its final generation. The serial program was run on the ∽10,000, ∽100,000, and ∽1x106 SNPs scenarios. As the speed of the CPU-based program is linear on the number of simulated SNPs, the resulting runtimes of 0.4, 3.9, and 38.7 seconds were then linearly rescaled to estimate runtimes for serial simulations with differing numbers of final SNPs. The two Intel processors have identical speeds on singlethreaded, serial tasks, which also allows for direct comparison between the two GPU results. Consumer GPUs like the 780M and 980 need to warm up from idle and load the CUDA context. So to obtain accurate runtimes on the GPU, *GO Fish* timings were done after 10 runs had finished and then the average of another 10 runs was taken for each data point. The *GO Fish* compacting rate was hand-optimized for each number of simulated SNPs, for each processor (Supplemental File 1).

Tools employing the single-locus Wright-Fisher framework are widely used in population genetics analyses to estimate selection coefficients and infer demography (see [11, 20-24] for examples). Often these tools employ either a numerically solved diffusion approximation, or even the simple analytical function, to generate the expected SFS of a given evolutionary scenario, which can then be used to calculate the likelihood producing an observed SFS (ref). The model parameters of the evolutionary scenario are then fit to the data by maximizing the composite likelihood (ref). With *GO Fish*, forward simulation can generate the expected spectra. To validate these expected spectra, the results of *GO Fish* simulations were compared against *δaδi* [15] for a complex evolutionary scenario involving a single population splitting into two, exponential growth, selection, and migration. (Figure 5) The spectra generated by each program are identical. Interestingly, the two programs also had essentially identical runtimes for this scenario and hardware. (Figure 5) In general, the relative compute time will vary depending on the simulation size for *GO Fish*, the grid size & time-step for *δaδi* [15], and the simulation scenario & hardware for both.

**Figure 5.**
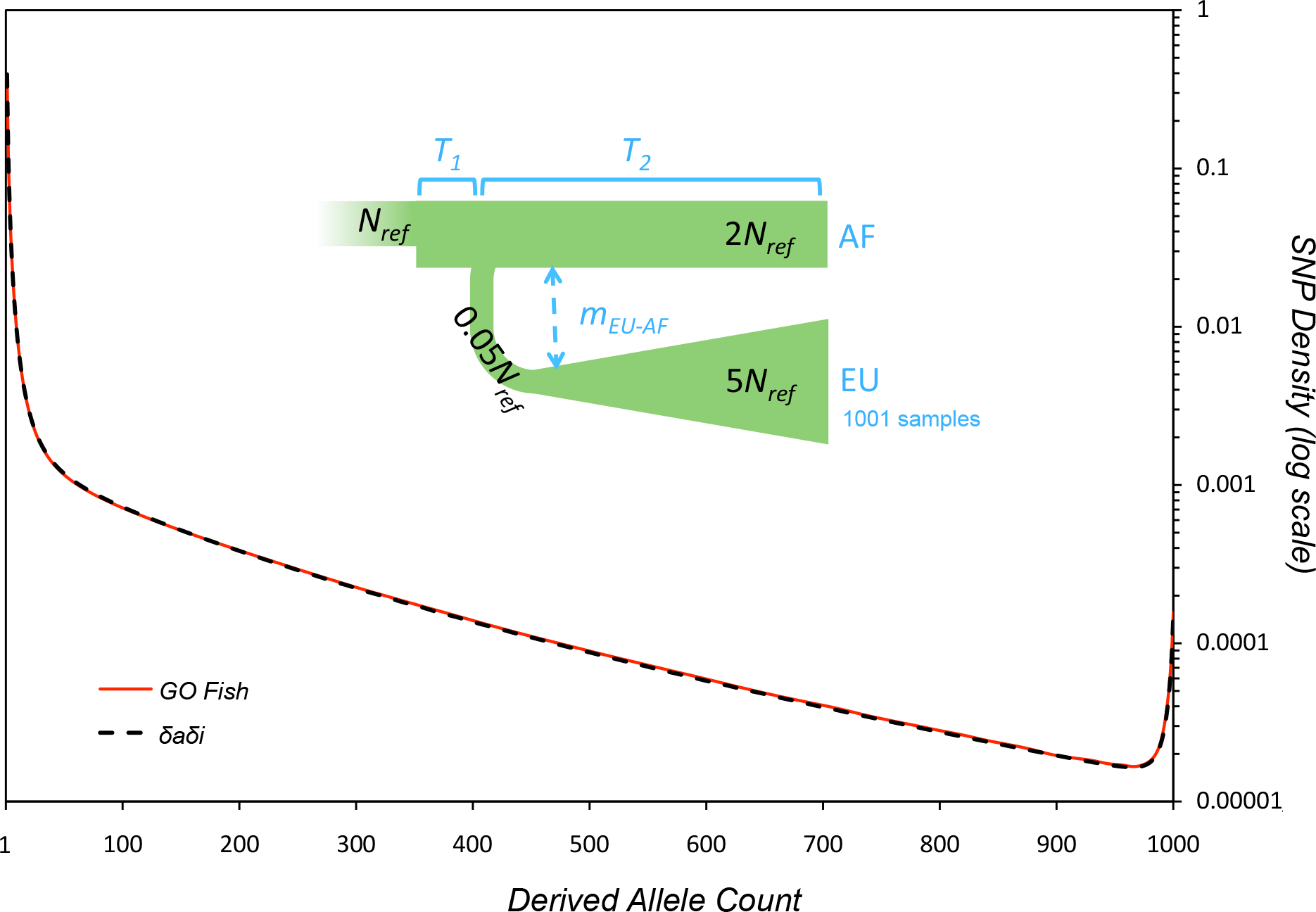
Validation of *GO Fish* simulation results against *δ aδ i*. A complex demographic scenario was chosen as a test case to compare the *GO Fish* simulation against an already established SFS method, *δaδi* [15]. The demographic model is from the YRI-CEU (AF-EU) *δaδi* example. Using *δaδi* [15] parameterization to describe the model, the ancestral population, in mutation-selection equilibrium, undergoes an immediate expansion from *N_ref_* to 2*N_ref_* individuals. After time *T_1_* (= 0.005) the population splits into two with a constant, equivalent migration, *mEU-AF* (= 1) between the now split populations. The second (EU) population undergoes a severe bottleneck of 0.05*N_ref_* when splitting from the first (AF) population, followed by exponential growth over time *T_2_* (= 0.045) to size 5*N_ref_*. The SFS (black dashed line) above is of weakly deleterious, co-dominant mutations (2*N_ref_s* = -2, *h* =0.5) where 1001 samples were taken of the EU population. The spectrum was then normalized by the number of segregating sites. The corresponding *GO Fish* parameters for the evolutionary scenario, given a mutation rate of 1x10^-9^ per site, 2x10^9^ sites, and an initial population size, *N_ref_*, of 10,000, are: *T_1_* = 0.005*2*N_ref_* = 100 generations, *T_2_* = 900 generations, *mEU-AF* = 1/(2*N_ref_*) = 0.00005, 2*Nrefs* = -4, *h* =0.5, and *F* = 0. As in *δaδi*, the population size/time can be scaled together and the simulation will generate the same normalized spectra [15]. Using the aforementioned parameters, a *GO Fish* simulation ends with ∽3x10^6^ mutations of which ∽560,000 are sampled in the SFS. The red line reporting *GO Fish* results is the average of 50 such simulations – the dispersion of those 50 simulations is reported in Figure S1. Each simulation run on the NVIDIA GeForce GTX 980 GPU took roughly the same time to generate the SFS as *δaδi* did (grid size = (110,120,130), time-step = 10^-3^) on the Intel Core i7 4771 CPU – just less than 0.7s.

For maximum-likelihood and Bayesian statistics as for parametric bootstraps and confidence intervals, hundreds, thousands, even tens of thousands of distinct parameter values may need to be simulated to yield the needed statistics for a given model. Multiplying this by the need to often consider multiple evolutionary models as well as nonparametric bootstrapping of the data, a single serial simulation run on a CPU taking only 18s, as in the simple simulation of ∽500,000 SNPs presented in Figure 4, can add up to hours, even days of compute time. Moreover, and in contrast to the approximating analytical or numerical solutions typically employed, simulating the expected SFS introduces random noise around the “true” SFS of the scenario being modeled. Figure S1 demonstrates how increasing the number of simulated SNPs increases the precision of the simulation – and therefore of the ensuing likelihood calculations. Simulating tens of millions of SNPs, wherein a single run on the CPU can take nearly half-an-hour, can be imperative to obtain a high-precision SFS needed for certain situations. Thus, the speed boost from parallelization on the GPU in calculating the underlying, expected SFS greatly enhances the practical utility of simulation for many current data analysis approaches. The speed and validation results demonstrate that, now with *GO Fish*, one can not only track allele trajectories in record time, but also generate SFS by using forward simulations in roughly the same time-frame as by solving diffusion equations. Just as importantly, *GO Fish* achieves the increase in performance without sacrificing flexibility in the evolutionary scenarios it is capable of simulating.

*GO Fish* can simulate mutations across multiple populations for comparative population genomics, with no limits to the number of populations allowed. Population size, migration rates, inbreeding, dominance, and mutation rate are all user-specifiable functions capable of varying over time and between different populations. Selection is likewise a user-specifiable function parameterized not only by generation and population, but also by allele frequency, allowing for the modeling of frequency-dependent selection as well as time-dependent and population-specific selection. By tuning the inbreeding and dominance parameters, *GO Fish* can simulate the full range of single-locus dynamics for both haploids and diploids with everything from outbred to inbred populations and overdominant to underdominant alleles. GPU-accelerated Wright-Fisher simulations thus provide extensive flexibility to model unique and complex demographic and selection scenarios beyond what many current site frequency spectrum analysis methods can employ.

Paired with a coalescent simulator, *GO Fish* can also accelerate the forward simulation component in forwards-backwards approaches (see [16, 25]). In addition, *GO Fish* is able to track the age of mutations in the simulation providing an estimate of the distribution of the allele ages, or even the age by frequency distribution, for mutations in an observed SFS. Further, the age of mutations is one element of a unique identifier for each mutation in the simulation, which allows the frequency trajectory of individual mutations to be tracked through time. This ability to sample ancestral states and then track the mutations throughout the simulation can be used to contrast the population frequencies of polymorphisms from ancient DNA with those present in modern populations for powerful population genetics analyses [26]. By accelerating the single-locus forward simulation on the GPU, *GO Fish* broadens the capabilities of SFS-analysis approaches in population genetic studies.

Across the field of population genetics and evolution, there exist a wide range of computationally intensive problems that could benefit from parallelization. The algorithms presented and discussed in Figure 2 represent a subset of the essential parallel algorithms, which more complex algorithms modify or build upon. Application of these parallel algorithms are already wide-ranging in bioinformatics: motif finding [27], global and local DNA and protein alignment [28-31], short read alignment and SNP calling [32, 33], haplotyping and the imputation of genotypes [34], analysis for genome-wide association studies [35, 36], and mapping phenotype to genotype and epistastic interactions across the genome [37, 38]. In molecular evolution, the basic algorithms underlying the building of phylogenetic trees and analyzing sequence divergence between species have likewise been GPU-accelerated [39, 40]. Further, there are parallel methods for general statistical and computational methods, like Markov Chain Monte Carlo and Bayesian analysis, useful in computational evolution and population genetics [41, 42].

Future work on the single-locus Wright-Fisher algorithm will include extending the parallel structure of *GO Fish* to allow for multiple alleles as well as multiple mutational events at a site, relaxing one of the key assumptions of the Poisson Random Field [4]. At present, neither running simulations with long divergence times between populations nor any scenario where the number of extant mutations in the simulation rises to too high a proportion of the total number of sites is theoretically consistent with the Poisson Random Field model underpinning the current version of *GO Fish*. Beyond *GO Fish*, solving Wright-Fisher diffusion equations in programs like *δaδi* [15] can likewise be sped up through parallelization on the GPU [43-46].

Unfortunately, while the effects of linkage and linked selection across the genome can be mitigated in analyses using a single-locus framework [15, 24, 47], these effects cannot be examined and measured whilst assuming independence amongst sites. Expanding from the study of independent loci to modeling the evolution of haplotypes and chromosomes, simulations with the coalescent framework or forward Wright-Fisher algorithm with linkage can also be accelerated on GPUs. The coalescent approach has already been shown to benefit from parallelization over multiple CPU cores (see [48]). While Montemuiño et al. achieved their speed boost by running multiple independent simulations concurrently, they noted that parallelizing the coalescent algorithm itself may also accelerate individual simulations over GPUs [48]. Likewise, multiple independent runs of the full forward simulation with linkage can be run concurrently over multiple cores and the individual runs might themselves be accelerated by parallelization of the forward algorithm. The forward simulation with linkage has many *embarrassingly parallel* steps, as well as those that can be refactored into one of the core parallel algorithms. The closely related *genetic algorithm*, used to solve difficult optimization problems, has already been parallelized and, under many conditions, greatly accelerated on GPUs [49-51]. However, not all algorithms will benefit from parallelization and execution on GPUs – the real world performance of any parallelized algorithm will depend on the details of the implementation [50, 51]. While the extent of the performance increase will vary from application to application, each of these represent key algorithms whose potential acceleration could provide huge benefits for the field [12, 13].

These potential benefits extend to lowering the cost barrier for students and researchers to run intensive computational analyses in population genetics. The *GO Fish* results demonstrate how powerful even an older, mobile GPU can be at executing parallel workloads, which means that *GO Fish* can be run on everything from GPUs in high-end compute clusters to a GPU in a personal laptop and still achieve a great speedup over traditional serial programs. A batch of single-locus Wright-Fisher simulations that might have taken a hundred CPU-hours or more to complete on a cluster can be done, with *GO Fish*, in an hour on a laptop. Moreover, graphics cards and massively parallel processors in general are evolving quickly. While this paper has focused on NVIDIA GPUs and CUDA, the capability to take advantage of the massive parallelization inherent in the Wright-Fisher algorithm is the key to accelerating the simulation and in the High Performance Computing market there are several avenues to achieve the performance gains presented here. For instance, OpenCL is another popular lowlevel language for parallel programming and can be used to program NVIDIA, AMD, Altera, Xilinx, and Intel solutions for massively parallel computation, which include GPUs, CPUs, and even Field Programmable Gate Arrays (FPGAs) [52-54]. The parallel algorithm of *GO Fish* can be applied to all of these tools. Whichever platform(s) or language(s) researchers choose to utilize, the future of computation in population genetics is massively parallel and exceedingly fast.

## Acknowledgements

The author would like to thank Nandita Garud, Heather Machado, Philipp Messer, Kathleen Nguyen, Sergey Nuzhdin, Peter Ralph, Kevin Thornton, and two anonymous reviewers for providing feedback and helpful suggestions to improve this paper.

## Appendix – Parallel Wright-Fisher Simulation Details

### Simulation Initialization

Simulations can be initialized in one of three ways: 1) a blank canvas, 2) from the results of a previous simulation, and 3) mutation-selection equilibrium. Starting a simulation as a blank canvas provides the most flexibility in what evolutionary state the simulation begins and thus any evolutionary scenario can be simulated from the beginning. However, as the simulation starts with no mutations present, a “burn-in” time is necessary to reach the point where the simulation of the scenario of interest can begin. The number of “burn-in” generations may be quite long, particularly to reach any kind of equilibrium state where selection, mutation, migration, and drift are all in balance and the number of mutations being fixed and lost is equal to the number of new mutations in the population(s). To save time, if a starting scenario is shared across multiple simulations, then the post-burn-in mutation array can be simulated beforehand, stored, and input as the initial mutation array for the next set of simulations.

Another way to jump start the simulation is by assuming all extant populations are in mutationselection balance at the beginning of the simulation. Under general mutation-selection equilibrium (MSE), the proportion of mutations at every frequency in the population can be calculated via numerical integration over a continuous frequency diffusion approximation (see [3]). While this constrains the starting evolutionary state to mutation-selection equilibrium, this allows one to then start simulating the selection and demographic scenario of interest immediately. Due to current limitations of the MSE model in *GO Fish*, the mutation-selection equilibrium scenario does not, as of yet, include migration from other populations or random fluctuations in selection intensity – nor can the code calculate the number of generations ago a mutation at frequency *x* is expected to have arisen at. Instead all mutations in the initial mutation array said to have arisen at time t = 0. The model is detailed below:

Using the glossary from Table 1, for any given population j at time t = 0:

*μ* = *μ*(*j*,0), s(*x*) = s(*j*,0,*x*), etc…

From Kimura p. 220-222 [3]:

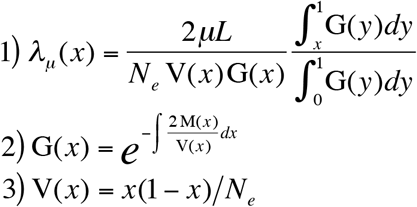

where *N_e_* = 2*N* (1 + *F*)

λ_μ_(*x*) is the expected (mean) number of mutations at a given frequency, *x*, in the population at mutation-selection equilibrium. V(*x*) and M(*x*) are the contribution of drift of selection respectively to the rate of change of a mutation’s frequency at frequency *y* in the population. Since this is an allele-based simulation, I use the equilibrium value of the effective number of chromosomes, *N_e_*, to account for inbreeding amongst *N* individuals.

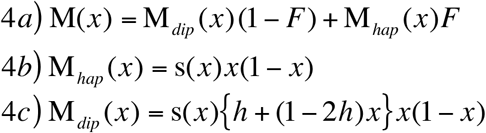

The total rate of frequency change is the average of the rate of change of the effective haploid proportion of the population and the effective diploid proportion of the population weighted by *F*.

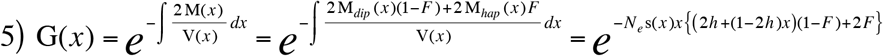

Substituting eq. 3 and 5 into eq. 1 yields:

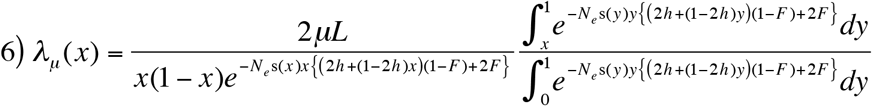

More familiar versions of eq. 6 can be derived by assuming neutrality or by assuming no frequency-dependent selection and either codominance or haploid/completely inbred individuals.

if s(*x*) = 0 ∀*x* ∈ (0,1) (neutral) → *λ_μ_* (*x*) = 2*μL x*
if s(*x*) = *s* ∀*x* ∈ (0,1) and (*h* = 0.5 or *F* = 1) → 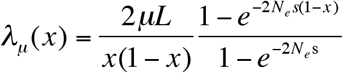
where if *h* = 0.5 (codominant) → *N_e_* = 2*N* (1+ *F*)
where if *F* = 1 (haploid) → *N_e_* = *N*

I approximate the integrals in eq. 6 using trapezoidal numerical integration and use the *scan* parallel algorithm implemented in CUB 1.6.4 [57] to accelerate the integration on the GPU*. λ_μ_(*x*) is the expected (mean) number of mutations. To determine the actual number of mutations at a given frequency, *x*, I generate random numbers from the Inverse Poisson distribution with mean λ_μ_(*x*) using the following procedure:

I. Random number generator Philox [58] generates a uniform random number between 0 and 1.
II. If λ_μ_(*x*) ≤ 6, then that uniform variable is fed into the exact Inverse Poisson CDF.
III. If λ_μ_(*x*) > 6, then a Normal approximation to the Poisson is used.

Adding all the new mutations at every frequency to the starting mutation array is an embarrassingly parallel problem. Thus, combined with the parallel numerical integration for the definite integral components of eq. 6, initializing the simulation at mutation-selection equilibrium is overall greatly accelerated on the GPU relative to serial algorithms on the CPU.

**\*An Aside About Numerical Precision, GPUs, and Numerical Integration:** For a bit of background, CPUs employ a Floating-point Processor Unit with 80-bits of precision for serial floating-point computation, which then quickly translates the result into double-precision (64-bit) for the CPU registers. Thus, CPU programs, including the serial Wright-Fisher simulation, are often written with double-precision performance in mind. In contrast, most consumer GPU applications are geared towards single-precision (32-bit) computation (e.g. graphics) and many consumer GPUs have relatively poor double-precision performance. More expensive, professional-grade workstation GPUs often have far better double-precision performance than their consumer counterparts. As the Wright-Fisher simulation does not actually require 64-bits of precision for its calculations, *GO Fish* has been written with 32-bits of precision computation in mind. This is even true of the MSE Integration step where the naturally pair-wise summation of parallel *scanning* mitigates the round-off error when performing large numbers of consecutive sums in 32-bit [59]. That said, the mutation frequencies stored in the simulation have only single-precision floating-point accuracy. Experiments using CPU serial Wright-Fisher simulations showed consistent results between storing mutation frequencies with 32-bits vs. 64-bits of precision.

#### Steps to *Calculate New Allele Frequencies*

Migration, selection, and drift determine the frequency of an allele in the next generation, *xt+1*, based on its current frequency, *xt*. Migration and selection are deterministic forces whereas drift introduces binomial random chance. While these three steps can, in principle, be done in any order, their order in the simulation is as follows:

I. Migration
II. Selection (with Inbreeding)
III. Drift (with Inbreeding)

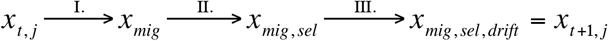

##### I. Migration

Using the glossary from Table 1, in population *j* at time *t*:

m(*k*) = m(*k, j, t*),
*x_t,k_* ≡ freq. of allele in pop. *k* at time *t*,
*x_mig_* = *x_mig, j_* ≡ freq. of allele in pop. *j* after migration,

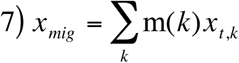

where 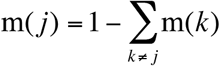

*GO Fish* uses a conservative model of migration [19]. The new allele frequency in population *j* is the average of the allele frequency in all the populations weighted by the migration rate from each population, to population *j*. And the migration rate is specified by the proportion of chromosomes from population *k* in population *j*.

##### II. Selection (with Inbreeding)

In population *j* at time *t*:

*x_mig_* = *x_mig, j_* ≡ freq. of allele after migration, *y_mig_* = 1 − *x_mig_*,
*x_mig, sel_* = *x_mig, sel_, j* ≡ freq. of allele after migration and selection,
*P_AA_*, *P_Aa_*, *P_aa_* ≡ frequency of genotype *AA, Aa*, and *aa*,
s(*x*) = s(*j, t, x*), h = h(*j*,*t*),
*w* = *w_j_* ≡ average pop. *j* fitness, *n* = *n_j_* ≡ average pop. *j* fitness of allele A

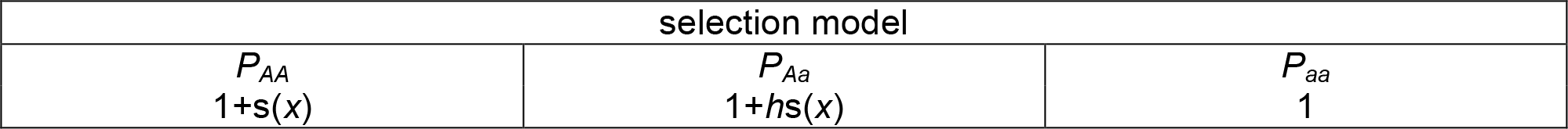

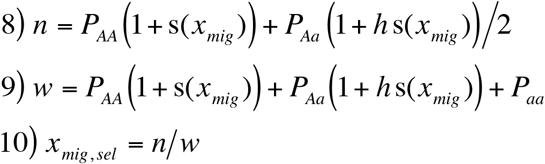

Like with M(*x*) in eq. 4, *w* and *n* are a weighted average of the effective haploid (inbred) and diploid (outbred) portions of the chromosome population. Diploid genotype frequencies assume random mating and Hardy-Weinberg equilibrium [60, 61].

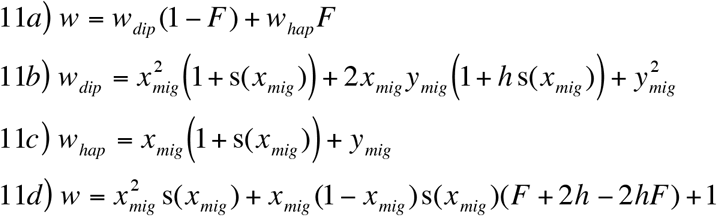

Following the same logic as above:

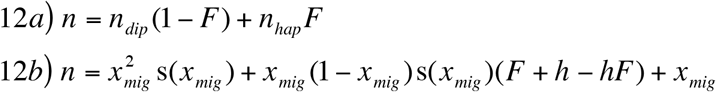

Substituting eq. 11*d* and 12*b* into eq. 10 yields:

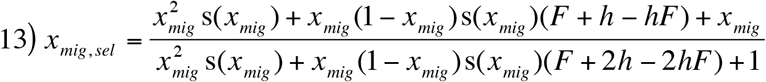

Again, like for eq. 6, more familiar forms of eq. 13 may be derived under certain assumptions such as neutrality, haploid/inbred individuals, and completely outbred diploids.

if s(*x_mig_*) = 0 ∀ *x* ∈ (0,1) (neutral) → *x_mig, sel_* = *x_mig_*
if *F* = 1 (haploid) → 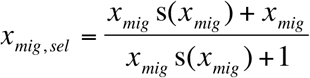
if *F* = 0 (diploid) → 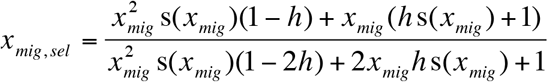

##### III. Drift (with Inbreeding)

For population *j* in generation *t*:

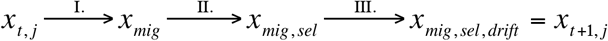

The variable *x_mig,sel_* represents the expected frequency of the allele in generation *t+1*. Drift is the random deviation of the actual frequency of the allele from this expectation. To determine the actual frequency of the allele in the next generation, *x_t+1,j_*, I generate random numbers from the Inverse Binomial distribution with mean *N_e_X_mig,sel_* and variance *N_e_X_mig,sel_*(1-*X_mig,sel_*) using the following procedure:

I. Random number generator Philox [58] generates a uniform random number between 0 and 1.
II. If *N_e_x_mig,sel_* ≤ 6, then that uniform variable is fed into the exact Inverse Poisson CDF as an approximation to the Binomial.
III. If *N_e_x_mig,sel_* > 6, then a Normal approximation to the Binomial is used.

As *N_e_* = 2*N*/(1+*F*), inbreeding affects drift as well as selection.

#### Adding New Mutations

Using the glossary from Table 1, for population *j* in generation *t*:

*μ* = *μ*(*j*,*t*), *N_e_* = 2N(*j*,*t*) (1 + *F*)
14) *λ_μ_* = *N_e_ μL*
starting frequency, *x* = 1/*N_e_*

The Poisson Random Field shares an important assumption with Watterson’s infinite sites model in that regardless of how many sites are currently polymorphic, mutations will never strike a currently polymorphic site and the number of monomorphic sites that a mutation can occur at is always the total number of sites, *L* [4, 62]. Eq. 14 defines the expected number of mutations in population *j* for generation *t+1*. The actual number of new mutations is drawn from the Inverse Poisson distribution using the same procedure detailed in *Simulation Initialization*. New mutations can be added to generation *t+1* in parallel and simultaneously with the new frequency calculations. Each new mutation is given a 4-part unique ID consisting of the thread and compute device that birthed it (if more than one graphics card is used) as well as the generation and population in which it first arose.

#### Compact

The general *compact* algorithm is outlined in Figure 2C). *GO Fish*’s version uses a more advanced variant to speed up compaction and lower its memory requirement as discussed in [18] and adapted from [63].

---

**Figure S1.**
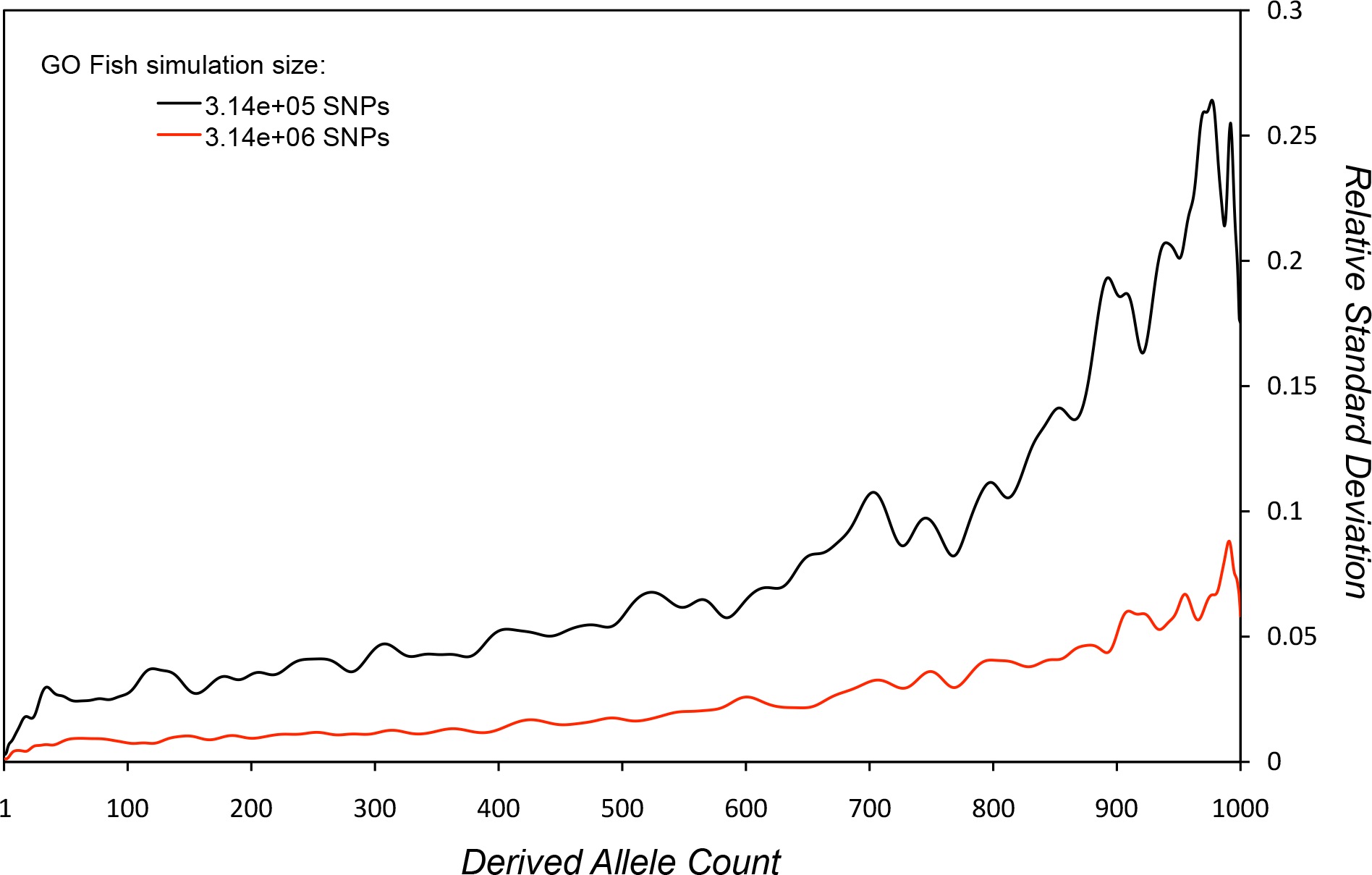
Dispersion of simulation runs over the SFS. Figure 5 shows the average SFS of 50 *GO Fish* simulation runs with ∽3x106 SNPs for a complex, multi-population evolutionary scenario – i.e. the mean estimated probability of seeing a SNP at each frequency in the sample of 1001 EU genomes given the evolutionary history. Dispersion of the results of these simulation runs for each allele count is measured by the relative standard deviation (σ/µ) and is displayed on the red line. The black line is the same evolutionary & sampling scenario but with a smaller simulation size, only ∽300,000 SNPs. As the mean probability of seeing a SNP at a frequency in the sample becomes more rare, the variance in the estimate of that probability across simulation runs increases. For example: at a derived allele count (DAC) of 1, the dispersion is ∽0.001/0.003 (red/black), whereas at a DAC of 1000, the dispersion is ∽0.058/0.175 (red/black). The factor of 10 separating the two simulation sizes engenders a roughly 3-fold difference in the dispersion (σ being the square root of the variance). Thus the expected precision with which an individual simulation run of 3x107 SNPs can ascertain the probability of seeing a SNP at a DAC of 1000 would be just under 2% for this evolutionary & sampling scenario. How much noise in the results of a simulation is acceptable for a particular application will determine the necessary simulation size.

## References

1. Fisher RA. (1930) The distribution of gene ratios for rare mutations. Proceedings of the Royal Society of Edinburgh 50: 205–220.

2. Wright S. (1938) The distribution of gene frequencies under irreversible mutation. Proc Natl Acad Sci U S A 24(7): 253.

3. Kimura M. (1964) Diffusion models in population genetics. J Appl Prob 1(2): 177–232.

4. Sawyer SA, Hartl DL. (1992) Population genetics of polymorphism and divergence. Genetics 132(4): 1161–1176.

5. Williamson S, Fledel-Alon A, Bustamante CD. (2004) Population genetics of polymorphism and divergence for diploid selection models with arbitrary dominance. Genetics 168(1): 463–475.

6. Lawrie DS, Messer PW, Hershberg R, Petrov DA. (2013) Strong purifying selection at synonymous sites in D. melanogaster. PLoS Genetics 9(5): e1003527.

7. Lawrie DS, Petrov DA. (2014) Comparative population genomics: Power and principles for the inference of functionality. Trends in Genetics 30(4): 133–139.

8. Hernandez RD. (2008) A flexible forward simulator for populations subject to selection and demography. Bioinformatics 24(23): 2786–2787.

9. Messer PW. (2013) SLiM: Simulating evolution with selection and linkage. Genetics 194(4): 1037–1039.

10. Thornton KR. (2014) A C++ template library for efficient forward-time population genetic simulation of large populations. Genetics 198(1): 157–166.

11. Ortega-Del Vecchyo D, Marsden CD, Lohmueller KE. (2016) PReFerSim: Fast simulation of demography and selection under the poisson random field model. Bioinformatics 32(22): 3516–3518.

12. Carvajal-Rodriguez A. (2010) Simulation of genes and genomes forward in time. Curr Genomics 11(1): 58–61.

13. Hoban S, Bertorelle G, Gaggiotti OE. (2012) Computer simulations: Tools for population and evolutionary genetics. Nature Reviews Genetics 13(2): 110–122.

14. Hudson RR. (2002) Generating samples under a wright-fisher neutral model of genetic variation. Bioinformatics 18(2): 337–338.

15. Gutenkunst RN, Hernandez RD, Williamson SH, Bustamante CD. (2009) Inferring the joint demographic history of multiple populations from multidimensional SNP frequency data. PLoS Genetics 5(10): e1000695.

16. Ewing G, Hermisson J. (2010) MSMS: A coalescent simulation program including recombination, demographic structure and selection at a single locus. Bioinformatics 26(16): 2064–2065.

17. Nickolls J, Buck I, Garland M, Skadron K. (2008) Scalable parallel programming with CUDA. Queue 6(2): 40–53.

18. Billeter M, Olsson O, Assarsson U. (2009) Efficient stream compaction on wide SIMD many-core architectures. ACM Proceedings of the conference on high performance graphics 2009: 159–166.

19. Nagylaki T. (1980) The strong-migration limit in geographically structured populations. J Math Biol 9(2): 101–114.

20. Koch E, Novembre J. (2017) A temporal perspective on the interplay of demography and selection on deleterious variation in humans. G3 (Bethesda) 7(3): 1027–1037.

21. Kim BY, Huber CD, Lohmueller KE. (2017) Inference of the distribution of selection coefficients for new nonsynonymous mutations using large samples. Genetics doi: genetics.116.197145 [pii].

22. Jackson BC, Campos JL, Haddrill PR, Charlesworth B, Zeng K. (2017) Variation in the intensity of selection on codon bias over time causes contrasting patterns of base composition evolution in drosophila. Genome Biology and Evolution 9(1): 102–123.

23. Garud NR, Messer PW, Buzbas EO, Petrov DA. (2015) Recent selective sweeps in north american drosophila melanogaster show signatures of soft sweeps. PLoS Genet 11(2): e1005004.

24. Machado HE, Lawrie DS, Petrov DA. (2017) Strong selection at the level of codon usage bias: Evidence against the li-bulmer model. bioRxiv : 106476.

25. Nakagome S, Alkorta-Aranburu G, Amato R, Howie B, Peter BM, et al. (2016) Estimating the ages of selection signals from different epochs in human history. Mol Biol Evol 33(3): 657–669.

26. Bank C, Ewing GB, Ferrer-Admettla A, Foll M, Jensen JD. (2014) Thinking too positive? revisiting current methods of population genetic selection inference. Trends in Genetics 30(12): 540–546.

27. Ganesan N, Chamberlain RD, Buhler J, Taufer M. (2010) Accelerating HMMER on GPUs by implementing hybrid data and task parallelism. ACM Proceedings of the First ACM International Conference on Bioinformatics and Computational Biology: 418–421.

28. Vouzis PD, Sahinidis NV. (2011) GPU-BLAST: Using graphics processors to accelerate protein sequence alignment. Bioinformatics 27(2): 182–188.

29. Zhao K, Chu X. (2014) G-BLASTN: Accelerating nucleotide alignment by graphics processors. Bioinformatics 30(10): 1384–1391.

30. Liu W, Schmidt B, Muller-Wittig W. (2011) CUDA-BLASTP: Accelerating BLASTP on CUDAenabled graphics hardware. IEEE/ACM Transactions on Computational Biology and Bioinformatics (TCBB) 8(6): 1678–1684.

31. Liu Y, Wirawan A, Schmidt B. (2013) CUDASW++ 3.0: Accelerating smith-waterman protein database search by coupling CPU and GPU SIMD instructions. BMC Bioinformatics 14: 117-2105–14-117.

32. Klus P, Lam S, Lyberg D, Cheung MS, Pullan G, et al. (2012) BarraCUDA - a fast short read sequence aligner using graphics processing units. BMC Res Notes 5: 27-0500–5-27.

33. Luo R, Wong T, Zhu J, Liu C, Zhu X, et al. (2013) SOAP3-dp: Fast, accurate and sensitive GPUbased short read aligner. PLoS ONE 8(5): e65632.

34. Chen GK, Wang K, Stram AH, Sobel EM, Lange K. (2012) Mendel-GPU: Haplotyping and genotype imputation on graphics processing units. Bioinformatics 28(22): 2979–2980.

35. Chen GK. (2012) A scalable and portable framework for massively parallel variable selection in genetic association studies. Bioinformatics 28(5): 719–720.

36. Song C, Chen GK, Millikan RC, Ambrosone CB, John EM, et al. (2013) A genome-wide scan for breast cancer risk haplotypes among african american women. PloS One 8(2): e57298.

37. Chen GK, Guo Y. (2013) Discovering epistasis in large scale genetic association studies by exploiting graphics cards. Frontiers in Genetics 4: 266.

38. Cebamanos L, Gray A, Stewart I, Tenesa A. (2014) Regional heritability advanced complex trait analysis for GPU and traditional parallel architectures. Bioinformatics 30(8): 1177–1179.

39. Suchard MA, Rambaut A. (2009) Many-core algorithms for statistical phylogenetics. Bioinformatics 25(11): 1370–1376.

40. Kubatko L, Shah P, Herbei R, Gilchrist MA. (2016) A codon model of nucleotide substitution with selection on synonymous codon usage. Mol Phylogenet Evol 94: 290–297.

41. Suchard MA, Wang Q, Chan C, Frelinger J, Cron A, et al. (2010) Understanding GPU programming for statistical computation: Studies in massively parallel massive mixtures. Journal of Computational and Graphical Statistics 19(2): 419–438.

42. Zhou C, Lang X, Wang Y, Zhu C. (2015) gPGA: GPU accelerated population genetics analyses. PloS One 10(8): e0135028.

43. Komatitsch D, Michéa D, Erlebacher G. (2009) Porting a high-order finite-element earthquake modeling application to NVIDIA graphics cards using CUDA. Journal of Parallel and Distributed Computing 69(5): 451–460.

44. Micikevicius P. (2009) 3D finite difference computation on GPUs using CUDA. ACM Proceedings of 2nd workshop on general purpose processing on graphics processing units: 79–84.

45. Tutkun B, Edis FO. (2012) A GPU application for high-order compact finite difference scheme. Comput Fluids 55: 29–35.

46. Lions J, Maday Y, Turinici G. (2001) A "parareal"in time discretization of PDE's. Comptes Rendus De L'Academie Des Sciences Series I Mathematics 332(7): 661–668.

47. Coffman AJ, Hsieh PH, Gravel S, Gutenkunst RN. (2016) Computationally efficient composite likelihood statistics for demographic inference. Mol Biol Evol 33(2): 591–593.

48. Montemuiño C, Espinosa A, Moure J, Vera-Rodríguez G, Ramos-Onsins S, et al. (2014) msPar: A parallel coalescent simulator. Springer Euro-Par 2013: Parallel Processing Workshops: 321–330.

49. Pospichal P, Jaros J, Schwarz J. (2010) Parallel genetic algorithm on the cuda architecture. In: Di Chio C, editors A, editor. Applications of Evolutionary Computation, EvoApplications 2010. Lecture Notes in Computer Science, vol 6024. Berlin, Heidelberg: Springer. pp. 442–451.

50. Hofmann J, Limmer S, Fey D. (2013) Performance investigations of genetic algorithms on graphics cards. Swarm and Evolutionary Computation 12: 33–47.

51. Limmer S, Fey D. (2016) Comparison of common parallel architectures for the execution of the island model and the global parallelization of evolutionary algorithms. Concurrency and Computation: Practice and Experience. doi: 10.1002/cpe.3797.

52. Stone JE, Gohara D, Shi G. (2010) OpenCL: A parallel programming standard for heterogeneous computing systems. Computing in Science & Engineering 12(1-3): 66–73.

53. Jha S, He B, Lu M, Cheng X, Huynh HP. (2015) Improving main memory hash joins on intel xeon phi processors: An experimental approach. Proceedings of the VLDB Endowment 8(6): 642–653.

54. Czajkowski TS, Aydonat U, Denisenko D, Freeman J, Kinsner M, et al. (2012) From OpenCL to high-performance hardware on FPGAs. IEEE 2012 22nd International Conference on Field Programmable Logic and Applications (FPL): 531–534.

55. Harris M. (2007) Optimizing parallel reduction in CUDA. NVIDIA Developer Technology 2(4). [ONLINE] https://docs.nvidia.com/cuda/samples/6_Advanced/reduction/doc/reduction.pdf.

56. Harris M, Sengupta S, Owens JD. (2007) Parallel prefix sum (scan) with CUDA. GPU Gems 3(39): 851–876.

57. Merrill D. (2016) CUB. v. 1.6.4 [ONLINE] https://nvlabs.github.io/cub/.

58. Salmon JK, Moraes M, Dror RO, Shaw DE. (2011) Parallel random numbers: As easy as 1, 2, 3. IEEE High Performance Computing, Networking, Storage and Analysis (SC), 2011 International Conference for: 1–12.

59. Higham NJ. (1993) The accuracy of floating point summation. SIAM Journal on Scientific Computing 14(4): 783–799.

60. Hardy GH. (1908) Mendelian proportions in a mixed population. Science 28(706): 49–50.

61. Weinberg W. (1908) Über den nachweis der vererbung beim menschen. Jahresh Wuertt Ver Vaterl Natkd 64: 369–382.

62. Watterson G. (1975) On the number of segregating sites in genetical models without recombination. Theor Popul Biol 7(2): 256–276.

63. Bakunas-Milanowski D, Rego V, Sang J, Yu C. (2015) A fast parallel selection algorithm on GPUs. IEEE 2015 International Conference on Computational Science and Computational Intelligence (CSCI): 609–614.

